# Long-range repression by ecdysone receptor on complex enhancers of the insulin receptor gene

**DOI:** 10.1101/2023.05.23.541945

**Authors:** Katie Thompson, Will Suber, Rachel Nicholas, David N. Arnosti

**Affiliations:** Department of Biochemistry and Molecular Biology; Department of Microbiology and Molecular Genetics, Michigan State University, East Lansing, Michigan

**Keywords:** Insulin, insulin receptor, Drosophila, enhancer, dFOXO, ecdysone, 20E, ecdysone receptor

## Abstract

The insulin signaling pathway is evolutionarily conserved throughout metazoans, playing key roles in development, growth, and metabolism. Misregulation of this pathway is associated with a multitude of disease states including diabetes, cancer, and neurodegeneration. Genome-wide association studies indicate that natural variants in putative intronic regulatory elements of the human insulin receptor gene (*INSR)* are associated with metabolic conditions, however, this gene’s transcriptional regulation remains incompletely studied. *INSR* is widely expressed throughout development and was previously described as a ‘housekeeping’ gene. Yet, there is abundant evidence that this gene is expressed in a cell-type specific manner, with dynamic regulation in response to environmental signals. The Drosophila insulin-like receptor gene (*InR*) is homologous to the human *INSR* gene and was previously shown to be regulated by multiple transcriptional elements located primarily within the introns of the gene. These elements were roughly defined in ∼1.5 kbp segments, but we lack an understanding of the potential detailed mechanisms of their regulation, as well as the integrative output of the battery of enhancers in the entire locus. Using luciferase assays, we characterized the substructure of these cis-regulatory elements in Drosophila S2 cells, focusing on regulation through the ecdysone receptor (EcR) and the dFOXO transcription factor. The direct action of EcR on Enhancer 2 reveals a bimodal form of regulation, with active repression in the absence of the ligand, and positive activation in the presence of 20E. By identifying the location of activators of this enhancer, we characterized a long-range of repression acting over at least 475 bp, similar to the action of long-range repressors found in the embryo. dFOXO and 20E have contrasting effects on some of the individual regulatory elements, and for the adjacent enhancers 2 and 3, their influence was/was not found to be additive, indicating that enhancer action on this locus can/cannot be characterized in part by additive models. Other characterized enhancers from within this locus exhibited “distributed” or “localized” modes of action, suggesting that predicting the joint functional output of multiple regulatory regions will require a deeper experimental characterization. The noncoding intronic regions of *InR* have demonstrated dynamic regulation of expression and cell type specificity. This complex transcriptional circuitry goes beyond the simple conception of a ‘housekeeping’ gene. Further studies are aimed at identifying how these elements work together in vivo to generate finely tuned expression in tissue- and temporal-specific manners, to provide a guide to understanding the impact of natural variation in this gene’s regulation, applicable to human genetic studies.

## Introduction

The conserved insulin signaling pathway plays a key role in development, reproduction, growth, and metabolism in metazoans (Brogiolo et al., 2001; Ebina et al., 1985; L. Petruzzelli et al., 1985; L. M. Petruzzelli et al., 1982). In mammals, insulin is released by the pancreas and binds to the insulin receptor (IR), a receptor kinase receptor that can be autophosphorylated, activating various metabolic pathways including the phosphatidylinositol 3-kinase (PI3K/AKT) pathway. Stimulation of this pathway is important for metabolic activity and furthermore involves the Ras-mitogen-activated protein kinase (MAPK) which is responsible for cell growth and development (Haeusler et al., 2018; Ullrich et al., 1985). FOXO is a key transcription factor that is phosphorylated in response to activation of the insulin signaling pathway. When FOXO is phosphorylated, it is excluded from the nucleus, abrogating the positive activity of this transcription factor on the insulin receptor gene (INSR), thus the FOXO-*INSR* relationship represents a negative feedback loop that may fine-tune levels of signaling (Puig et al., 2003; Puig & Tjian, 2005).

Signaling through the insulin receptor has been associated with a number of conditions relevant to human health. Rare mutations in the insulin receptor protein coding sequence can give rise to severe growth defects and insulin resistance (Accili et al., 1989; Ardon et al., 2014), while changes in the expression and regulation of *INSR* can affect overall signaling and has been associated with diabetes, cancer, and neurodegenerative diseases. (Craft et al., 2013; Czech, 2017; Frölich et al., 1998; Vella et al., 2018; Westermeier et al., 2016). In Type II diabetes, insulin binds to the insulin receptor but there is a failure to activate the insulin signaling cascade, resulting in a loss of glucose transport into cells and ultimately high blood glucose levels (Czech, 2017; Westermeier et al., 2016). The insulin receptor gene comprises 22 exons and 21 introns; alternative splicing at exon 11 results in two isoforms of the protein, IR-B, and IR-A. IR-B is the dominant, mature isoform while IR-A is found predominantly in fetal cells and cancer cells. The overexpression of IR-A increases the isoform ratio IR-A/IR-B up to 20-fold in some cancers and allows cancer cells to respond to insulin and insulin-like growth factors (Belfiore & Malaguarnera, 2011; Czech, 2017; Flannery et al., 2016; Vella et al., 2018). The insulin signaling pathway is also critical in the process of aging and Alzheimer’s disease (AD).; insulin regulates brain glucose metabolism in the brain, but in AD there is often reduced IR expression and tyrosine kinase signaling, resulting in defects in neuronal activity and cognitive function (Craft et al., 2013; Frölich et al., 1998). The insulin receptor thus provides a potential target for therapies for these diseases, including interventions that may impact binding affinity of the insulin to the receptor, by increasing expression cell surface receptors, or by prevention of ubiquitination and degradation of the receptors (Zhang et al., 2022). We lack a general understanding of how *INSR* expression levels impact physiology and disease and its potential as a target for therapeutic purposes. *INSR* has been previously described as a ‘housekeeping’ gene because of its broad expression, however, there is evidence of cell type specificity and cis-regulatory elements (Leal et al., 1992; Payankaulam et al., 2019; Wei et al., 2016).

The human *INSR* and the *InR* in Drosophila are homologous genes that are broadly expressed. The genes include large introns that have been demonstrated to contain chromatin features predicted to possess enhancer function or have been demonstrated to direct transcription (Wei et al. 2016; Payankaulam et al. 2019). In insects, the insulin signaling pathway has also been found in most tissues and shown to be crucial for embryonic development, neuronal function, and organ growth (Brogiolo et al., 2001; L. Petruzzelli et al., 1985; Song et al., 2003). Insulin-like peptides, dilps, are released by the prothoracic gland, PG, and bind to the insulin-like receptor, InR, leading to a similar insulin signaling cascade as seen in mammals (Brogiolo et al., 2001). In Drosophila as in mammals, the insulin signaling pathway responds to the environment and signaling inputs to regulate gene expression of the *insulin receptor*. Signaling involves the action of dFOXO, the Drosophila homolog of FOXO (Puig et al., 2003; Puig & Tjian, 2005). A central steroid hormone in Drosophila is 20-hydroxyecdysone (20E), which binds to the ecdysone receptor (EcR) and triggers activation of diverse genes that are critical for development. Surges of ecdysone during different developmental stages engage batteries of genes that can be distinguished between early/late genes. (Ashburner et al., 1974; Burtis et al., 1990; Segraves & Hogness, 1990) EcR has been shown to directly bind throughout the entire Drosophila genome including, usually within 10 kbp of 20E-regulated genes in a tissue-specific manner. The pervasive effects of ecdysone signaling can be attributed to the many transcription factors whose genes are targeted by this pathway (Gauhar et al., 2009). Ecdysone has been shown to regulate *InR* expression, but there is a lack of understanding of the direct action of EcR on the *InR* locus (Gauhar et al., 2009; Koelle et al., 1991).

We previously identified discrete regulatory regions within the intronic regions of the *InR* locus in Drosophila. Utilizing luciferase assays in cultured *Drosophila melanogaster* S2 and Kc cells, the ∼40 kbp of intronic regions were assessed as 25 subfragments of ∼1.5 kbp each (1-25) including those responsive to dFOXO or 20E treatment (Fig. 1). The active elements were then further mutagenized by specific serial deletions to identify parts of the enhancer that were necessary for activity. Some enhancers functioned in a cell type-specific manner, with preferential action in S2 vs. Kc cells (Wei et al., 2016).

**Figure 1:**
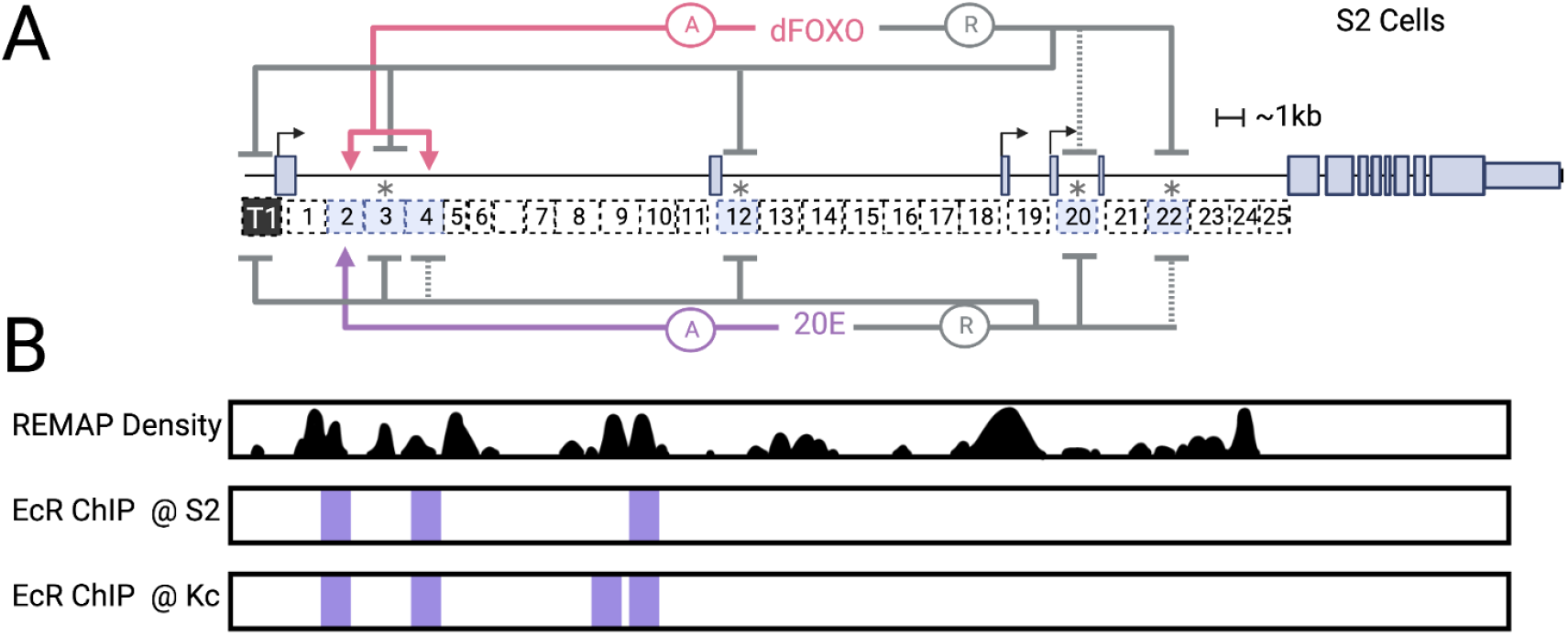
Regulatory elements of the Drosophila InR locus in S2 Cell Culture. **A**. Previous findings with luciferase reporter assays in S2 cell culture identification of enhancers on the *InR* locus. (Wei et al., 2016) *intrinsic activators **B**. REMAP 2022 Density showing regulatory regions throughout *InR* locus (Hammal et al., 2022). REMAP is a large-scale integrative analysis of transcriptional regulators that catalogs the results of ChIP-seq, ChIP-exo, and DAP-seq. EcR ChIP-seq peaks in S2 and Kc cells from the UCSC Genome Browser (Kent et al., 2002). (Recreated with Adobe Illustrator)

In this study, we sought to learn about the molecular organization of these putative *InR* enhancer elements by testing for sufficiency of small regulatory blocks, and not merely necessity, as in our earlier work. Our analysis of sub-elements 300 bp and 600 bp in size revealed diverse activities; in some cases, the action of the larger regulatory region can be summarized as the activity of one or two individual small elements, but in other cases, activity was dependent on much larger fragments, suggesting the cooperative action of factors binding across these elements. Particularly striking was the identification of a long-range repressor action mediated by the ecdysone receptor, a protein that has largely been studied in the context of promoter-proximal activity.

## Results

### Distinct actions of promoter-proximal enhancers 2, 3, and 4

Our previous studies demonstrated that there are active elements, designated enhancers 2-4, located within 2.3 kbp of the main transcriptional start site that shows intrinsic action in Kc and S2 cells and/or are affected by signaling inputs dFOXO and 20E. This segment of regulatory elements is interesting because it appears to integrate signaling inputs in a diametrically opposed fashion, with enhancer 2 stimulated by treatments that appear to reduce enhancer 3 activity. We carried out a fine-structure analysis to identify elements sufficient for regulation, with the goal of understanding possible independent or “integrative” activities (Wei et al., 2016). All of the constructs were tested in the context of the T1 promoter, which is the major promoter utilized for *InR* in S2 cells and other cell types. This promoter region, which extends from -900 to +250, contains more than the core basal promoter and may contribute to regulated expression of the gene. Indeed, although this element has low intrinsic activity in S2 cells, we found that activity of the T1 promoter construct alone was significantly affected by overexpression of dFOXO and 20E treatment (Fig 1A, 3A). The observed repression effect may be an indirect one, as T1 lacks known binding of EcR from ChIP data, and does not have canonical dFOXO binding motifs.

Enhancer 2 demonstrates low intrinsic activity in S2 cells, but is strongly upregulated upon overexpression with dFOXO or treatment with 20E. We further subdivided this element into 300 bp and 600 bp elements to test for sufficiency of action (Fig. 2, 3B). The treatment with dFOXO revealed significant activation centered on the 5’-most 2ab/2a fragment. In contrast, for 20E treatment, the more centrally-located 2bc/2cd/2c fragments were induced by this treatment, while lacking significant activity on their own in the absence of 20E. Our previous serial deletions of the full-length element demonstrated that removal of 2a reduced dFOXO-stimulated activity, and removal of 2c increased the inherent activity of enhancer consistent with our findings that dFOXO acts through 2a, while 2c has a repression and activation function, that is sufficient and necessary for response to 20E (Wei et al., 2016).

**Figure 2:**
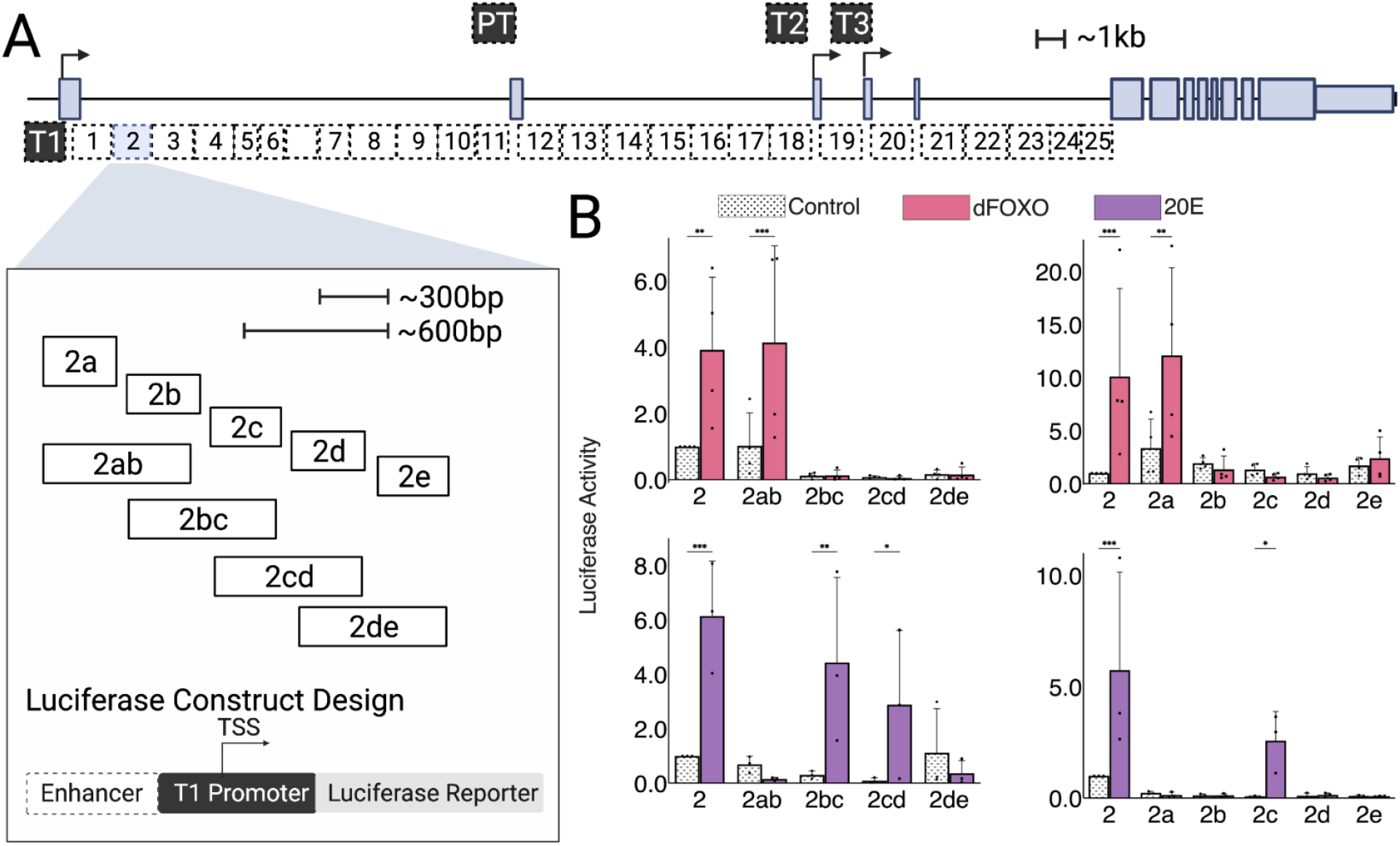
Luciferase reporter analysis of *InR* regulatory region enhancer 2 has separable dFOXO and 20E elements. **A**. *InR* intronic region broken into 25 ∼1.5kb fragments (1-25). Enhancer 2 was cloned into smaller 300 bp and 600 bp fragments (2a-2e,2ab-2de). Each enhancer fragment was cloned into a luciferase reporter upstream of the *InR* T1 basal promoter. **B**. Enhancer 2 300 bp and 600 bp fragments treated with dFOXO or 20E. Normalized to the full enhancer control.

**Figure 3:**
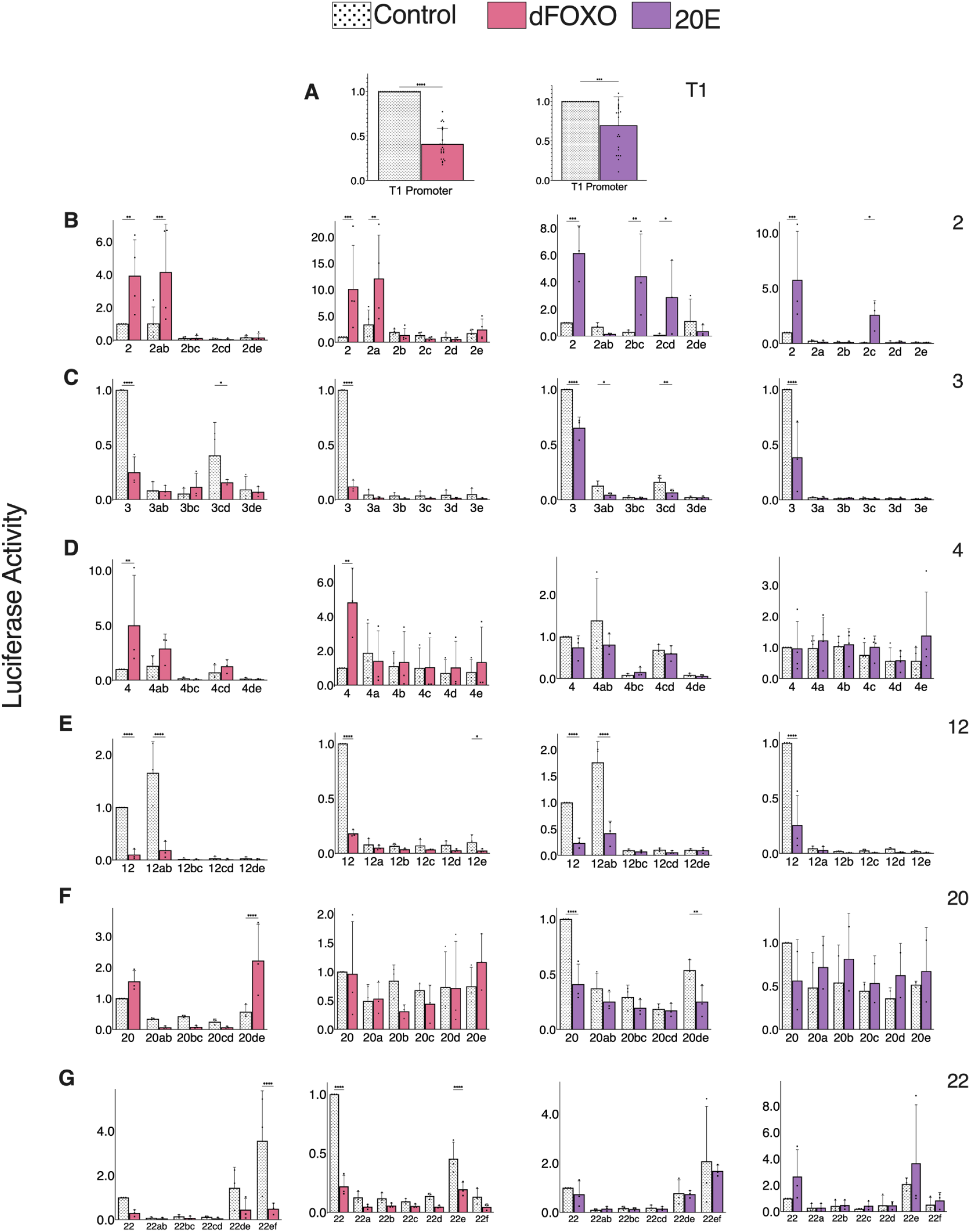
Luciferase reporter analysis of *InR* regulatory enhancers 300 bp and 600 bp fragments. **A**. T1 Promoter response to dFOXO overexpression and treatment with 20E. **B-G:** Enhancers 300 bp and 600 bp fragments treated with dFOXO or 20E. Normalized to the full enhancer control.

Enhancer 3 has a high intrinsic activity in S2 cells and is repressed by both dFOXO overexpression and 20E treatment. The 300 bp and 600 bp fragments derived from enhancer 3 showed overall low activity, suggesting that none of these elements are large enough to recapitulate intrinsic activity (Fig. 3C). Weak intrinsic activity was observed from 3cd, and possibly 3ab, but the total of these did not equal the intact element; it is likely that multiple binding sites for transcriptional activators distributed throughout the element are needed for function. Our previous serial deletions of enhancer 3 indicated that removal of either 3a, 3c, or 3d severely compromises output of enhancer 3, thus a “distributed” architecture of activators appears to be characteristic of this element. The negative effects of dFOXO and 20E may reflect indirect effects mediated through the T1 promoter region, which would reduce potential output of enhancer 3. Alternatively, an indirect genetic cascade may impinge on the enhancer itself. Unlike enhancer 2, it is not clear if these effects are impacting the same or distinct sub-elements of enhancer 3.

Enhancer 4 has low intrinsic activity and is stimulated by dFOXO overexpression, but not by 20E treatment. The 300 bp subfragments lacked any dFOXO response, but the 600 bp 4ab fragment appeared to exhibit stimulation by dFOXO. (Fig. 3D) Interestingly, the 4bc and 4de fragments show activity lower than the T1 promoter alone, suggesting the presence of repressors within these elements. The lack of intrinsic activity as well as dFOXO response by any of the 300 bp elements indicates that functionality is distributed over larger elements in enhancer 4. This enhancer was the only dFOXO-inducible element that exhibited cell-type activity (no response in Kc cells) indicating that dFOXO responsiveness may be enabled by distinctive sets of factors present on these enhancers some of which are restricted to S2 cells.

### EcR long-range repressor activity

EcR regulation of InR may include direct and indirect effects; previous ChIP experiments have identified binding within regions enhancer 2, 4, and 10 (Fig. 1B) Significantly, one peak overlaps the 2c portion of enhancer 2. We noted that there is an evolutionarily conserved region of this element that includes a highly conserved EcR motif (Fig. 4B). To test if this EcR binding motif mediated regulation by 20E we deleted the predicted binding sequence from the 300 bp 2c element (ΔEcR) and compared the activity of the mutant element to the wild-type (Fig. 4A). The 2c ΔEcR mutant element was inactive and not induced when treated with 20E, indicating that the motif is essential for regulation (Fig. 4C). Interestingly, we found that the same mutation introduced into the full-length enhancer 2 caused basal activity to significantly increase, to approximately the same level as that of the 20E-stimulated wild-type enhancer (Fig. 4C). The treatment of this element with 20E reduced activity, perhaps via inhibitory effects on the basal promoter (Fig. 3A). dFOXO overexpression stimulated both wild-type enhancer 2 as well the enhancer 2 ΔEcR mutant, although the latter had a higher baseline, and further stimulated by dFOXO, indicating that on the wild-type element, EcR repression reduces the potential activity of dFOXO (Fig. 4C). Treating with both 20E and dFOXO led to net induction of both wild-type and ΔEcR mutant forms of enhancer 2 (Fig. 4C).

**Figure 4:**
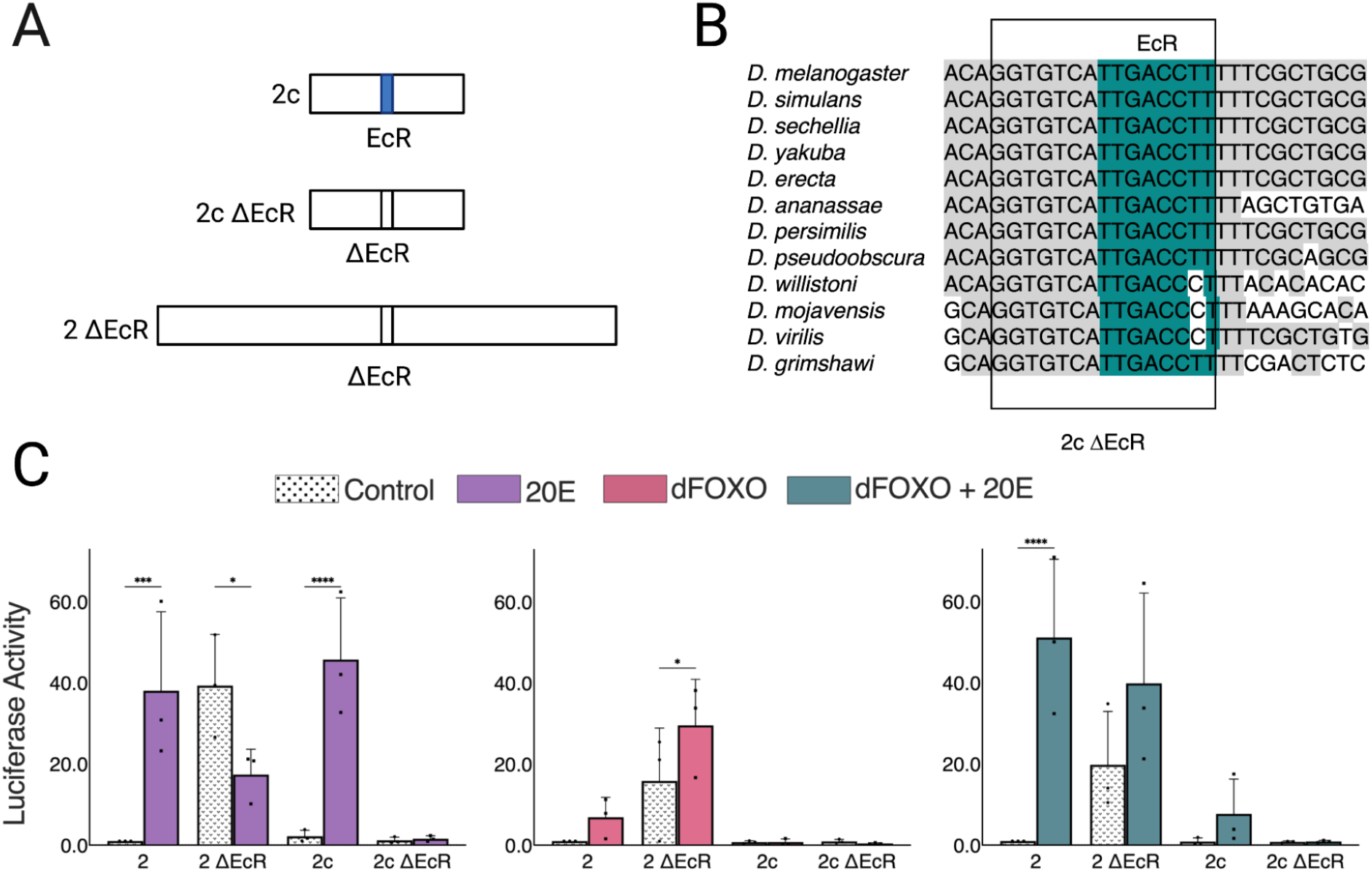
Long-distance ecdysone receptor repression of dFOXO on enhancer 2. **A**. Site-directed mutagenesis to delete EcR binding site localized at enhancer 2c. Deletions were made on the full enhancer 2 and 300bp fragment 2c. **B**. Evolutionarily conserved EcR binding site throughout 12 Drosophila species. **C**. Deletion of EcR on 2c and 2 full treated with 20E, dFOXO, or dFOXO + 20E.

These results suggest that in the wild-type enhancer, repression mediated by the EcR site curtails the activity of distal regulation by activators located within region 2a. The action of EcR over a distance of at least 475 bp is consistent with the activity of long-range repressors such as Hairy, which have the ability to interfere with distant activators by chromatin modifications that extend over a span of hundreds of bp (Kok et al., 2015; Li & Arnosti, 2011). Our previous serial deletions, taking out large blocks of sequence, are consistent with the distal action of a repressor in 2c and inherent activation by 2a. EcR thus appears to have a bimodal activity; such a repressor-to-activator switch of this factor has been previously observed and is consistent with the conserved mechanisms of other hormone and steroid receptors (Cherbas et al., 1991).

### Combinatorial action of enhancers 2 and 3

The enhancers we defined were arbitrarily divided into 1.5 kbp segments for the sake of analysis, but these sequences are contiguous in the endogenous gene. Regulatory elements located within them may interact or function autonomously to control *InR* promoter activity. Having looked at their independent actions, we combined enhancer 2 and enhancer 3 in their native configuration (Fig. 5A). Here, we also tested the variant enhancer 2 lacking the EcR motif. Overall activity of wild-type 2+3 was similar to 3 alone, which would be consistent with an additive action (Fig. 5B), while in the presence of 20E, the stimulated activity was greater. There may be elements in these two enhancers that synergize to produce greater than additive outputs in the absence of EcR repression. For the 2+3 ΔEcR construct, the activity has a higher baseline activity than wild-type 2+3, presumably reflecting the removal of EcR repression action, and here the output appears to be the sum of the potential for enhancer 2 alone with removal of EcR repression (the 20E treatment value) and inherent activity of 3. 20E treatment of this combination leaves output unchanged; the somewhat negative effect of 20E on 3 alone appears to be mitigated. Overall, this combination of the 2+3 regulatory region appears to produce results that are aligned with an additive model for these enhancers. However, the artificial placement within a reporter plasmid system may bias these results, as enhancer 3 is positioned in a non physiological promoter proximal position.

**Figure 5:**
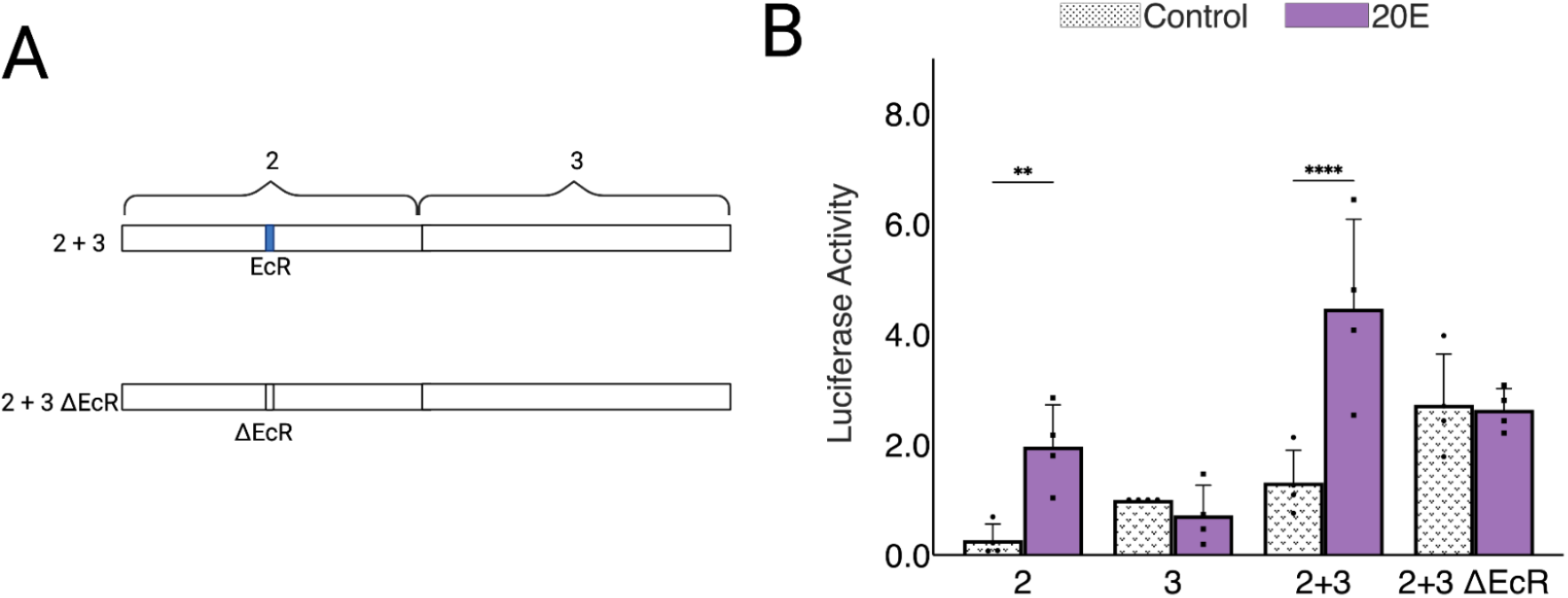
Combinatorial mechanism of enhancer 2 + enhancer 3 with deletion ecdysone receptor binding site. **A**. Site-directed mutagenesis to delete EcR binding site on enhancer 2 + enhancer 3. **B**. Deletion of EcR on enhancer 2 + enhancer 3 treated with 20E. Normalized to Full enhancer 3 control.

### Enhancers 12, 20, and 22

Enhancer 12 possesses a relatively high intrinsic activity, similar to that of enhancer 3. When enhancer 12 is subdivided into the 300 bp or 600 bp fragments, we find that the 3ab element is highly active, even somewhat higher than the parental element. This increase in activity may reflect loss of a repressive activity in more 3’ portions of the 1.5 kbp element (Fig. 3E). The closer proximity to the T1 promoter in the luciferase vector may also play a role. Interestingly, none of the 300 bp subelements exhibited much activity, suggesting that dividing a-b in half separates cooperating elements. The response of the active 3ab element to overexpression of dFOXO or treatment with 20E recapitulates the effect seen on the intact 1.5 kbp element, suggesting that transcriptional impacts are mediated by the 3ab portion. The reduction of expression by both of these treatments is much greater than the less-than-twofold effect observed on the T1 promoter alone, suggesting that the enhancer itself mediates this effect. These results are consistent with our previous internal deletions of enhancer 12, which showed that 12a is necessary for activation, and 12c-d mediates some sort of repression (Wei et al., 2016).

Enhancer 20 is also intrinsically active and is repressed by 20E, while the impact of dFOXO overexpression is ambiguous. In our earlier study, it appeared to be unaffected or repressed, though this effect was not statistically significant. Here we see an apparent induction, but this was similarly not statistically significant. Notably, the 600 bp 20de fragment was significantly upregulated by dFOXO overexpression, but neither the 300 bp 20d nor 20e fragments alone exhibited this effect. 20E repression was significant for both the parental 1.5 kbp enhancer as well as the 20de fragment. 300 bp fragments failed to show this effect. (Fig. 3F). Unlike enhancer 2, which has distinct elements for dFOXO and 20E regulation, for enhancer 20, the responses are colocalized on 20de. The previous serial deletions within this 1.5 kbp element showed that 20e, and 20b to a lesser extent, are necessary for intrinsic activity in S2 cells (Wei et al. 2016). In these previous studies, removal of 20a or 20b endowed the previously somewhat unresponsive enhancer with a potent induction upon dFOXO overexpression, suggesting that a repressive function within the 5’ portion of enhancer 20 limits dFOXO potential regulation. 20E regulation, in contrast, was uniformly inhibitory for these mutants, regardless of starting activity. In contrast, in our examination of subfragments, for the 300 bp elements, none were reproducibly regulated by 20E treatment; among 600 bp elements, only 20de (which lacks the inhibitory a-b region) was strongly downregulated by 20E. Overall, these experiments underscore the distinction between dFOXO and 20E regulation on this single enhancer element.

Regulation of enhancer 22 is characterized by activity solely in the 3’ most fragments of both 300 bp and 600 bp, which is suppressed by overexpression of dFOXO, with no significant response to treatment of 20E. (Fig. 3G). Our previous serial deletions of enhancer 22 are consistent with these results, showing the importance of the terminal 600 bp for activity, with a possible repressor activity in the 5’ portion (here, our minimal elements would not reveal a repressor in 22a, since it would be inactive by itself) (Wei et al., 2016). Overall, this detailed characterization of the minimal elements sufficient for activity supports our earlier conclusions that dFOXO and 20E have overlapping but separable actions on most enhancers and that the enhancers appear to be integrating contrasting regulatory information i.e. activation or repression by these signals. The detailed dissection of these elements also reveals the long-range effect of EcR repression within the enhancer 2 region, which does not appear to extend more broadly across the enhancer 2-3 region.

### Evolutionary Conservation

Having defined the functional elements of *InR* necessary and sufficient for activity in S2 cells, it is interesting to speculate on the conservation of such cis-regulatory circuitry. Within Drosophila, the protein coding exons of *InR* are highly conserved throughout evolution, as expected, while the intronic regions are less conserved (Fig. 6A). Enhancer sequences are difficult to identify with sequence analysis alone, although these sequences are evolutionarily constrained and can be enriched in functional variants relevant to complex traits and disease (Hindorff et al., 2009). However, by combining information from reporter assays, evolutionary conservation, and chromatin marks, we can obtain a clearer picture of *InR* regulatory elements. Considering the cluster of promoter-proximal enhancers (2-4), enhancer 2 is more highly conserved than enhancers 3 and 4, perhaps underscoring the importance of the direct EcR regulatory switch in regulating *InR* (Fig. 6B).

**Figure 6:**
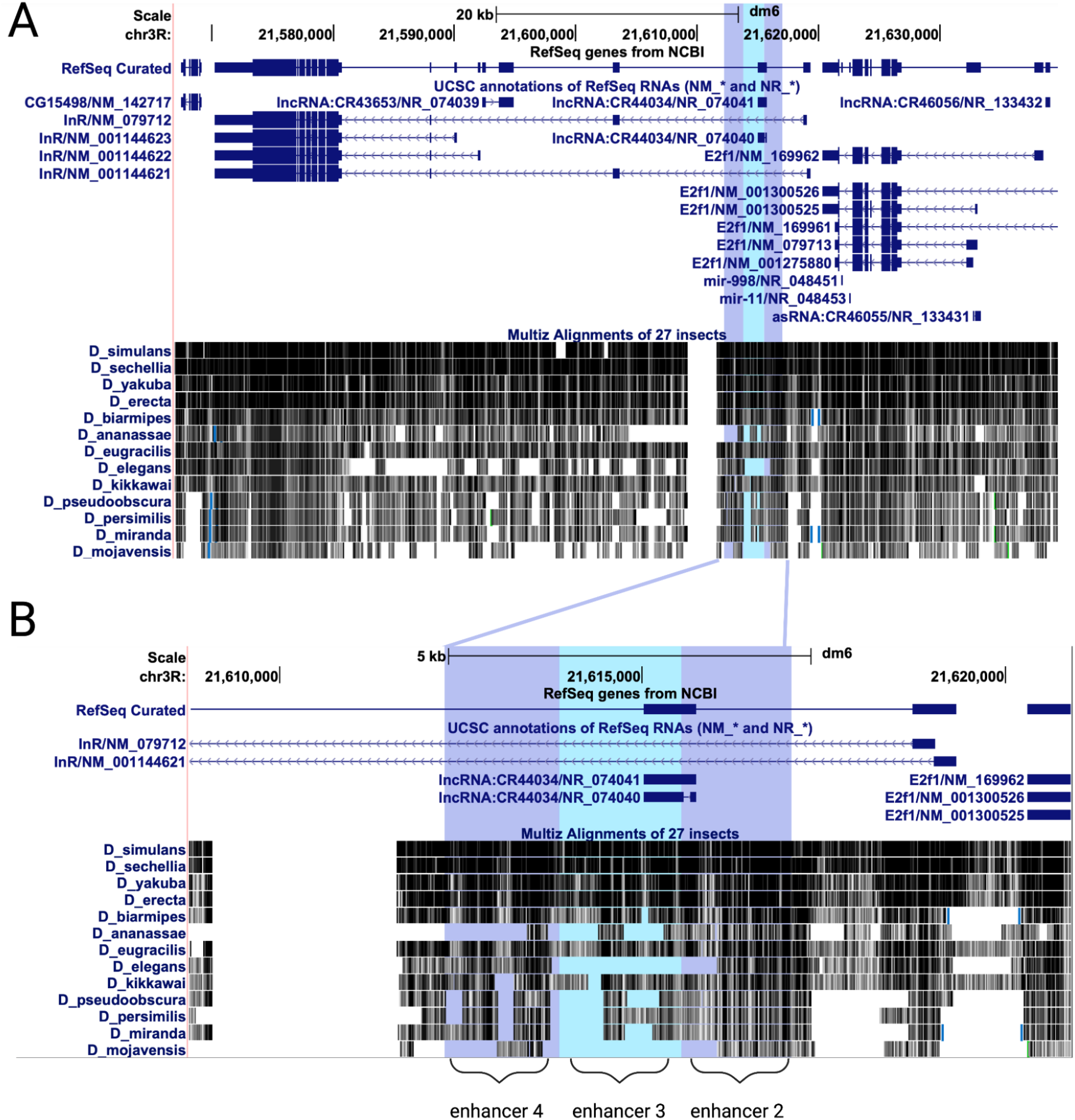
Evolutionary conservation of the *InR* locus. **A**. Conservation of the entire InR locus in 13 Drosophila species from the UCSC Genome Browser (Kent et al., 2002). **B**. Zoomed in view of the integrated locus of enhancers 2, 3, and 4.

## Discussion

EcR has been shown to regulate hundreds of loci in Drosophila, although many genes controlled by 20E represent secondary or tertiary effects of the hormone. Direct regulation by EcR has been demonstrated in a number of cases. For instance, EcR binds to promoter-proximal regions of *hsp27* and *fbp1* and greatly stimulates their expression upon exposure to 20E (Antoniewski et al., 1996). This on/off effect represents an important mechanism for driving cell- and developmental-specific gene expression. Notably, about one half of identified EcR interaction sites identified in Kc cells did not result in changes to gene expression of nearby genes, but many of these “unresponsive” sites showed evidence of 20E regulation in other cell types (Gauhar et al., 2009). Therefore, EcR may require additional factors in some instances to enable hormonal regulation. We see evidence of this phenomenon on *InR* as well. ChIP-seq identified EcR binding sites within the insulin-like receptor gene in S2 cells on enhancers 2, 4, and 10 (Fig 1B). The only enhancer to have an ecdysone response in S2 cells however was enhancer 2. Enhancer 4 exhibited no significant response to ecdysone, and enhancer 10 had a response only in Kc cells (Wei et al., 2016). The other sites bound by EcR may represent enhancers active in other cell types. Thus, defining the functional significance of EcR binding thus requires direct experimentation. By making use of transfected reporters in Drosophila cell culture, we uncovered novel aspects of *InR* regulation, in particular the direct regulation by EcR, and the range of repression mediated by this factor when not liganded to 20E. Previous studies demonstrated that EcR is found not only at the promoter, but also in distal locations, such as those on early genes such as *E75, E74*, and *Broad* (Bernardo et al., 2014). On these genes, stimulation by 20E induces the formation of enhancer-promoter loops that reflect promoter activation. These genes are, however, not as widely expressed as *InR is*, and these studies did not explore whether EcR was functioning as an active repressor in the absence of hormone, although previous studies have found that EcR can indeed associate with corepressors ((Brodu et al., 1999; Cherbas & Cherbas, 1981). Thus, the question arises whether EcR mediates a binary on/off activity on *InR* as well. Our findings indicate that within the *InR* intronic region comprising enhancer 2, EcR locally represses activators., Enhancer 2 repression does not appear to affect the activity of enhancer 3 when the enhancers are combined (Fig 5B). Thus, a locally-acting binary operation by EcR, embedded in a larger field of regulatory elements, may provide *InR* with a steadier output, suitable for this broadly expressed gene. Such localized repression function may also explain how repressors and corepressors participate in so-called “soft repression” (Mitra et al., 2021).

FOXO regulation of *InR* has been previously identified as an important feedback mechanism that may potentiate insulin signaling in Drosophila as well as mammals (Puig & Tjian, 2005). Previous studies in Drosophila have suggested that dFOXO may interact with the gene through internal promoters; however, the experimental evidence for this is weak (Orengo et al., 2017) There is limited information regarding *in vivo* binding by this transcription factor in Drosophila. A study exploring dFOXO binding in the adult female identified ∼700 sites, dFOXO was bound to the coding region of *InR* in adult females and previously shown to bind the T1 promoter in S2 cells (Alic et al., 2011; Puig et al., 2003; Puig & Tjian, 2005). Our previous ChIP identified dFOXO binding activity within the *InR* gene in S2 cells. Yet, the most prominent peaks on region 10 did not correspond to dFOXO responsiveness, and binding at enhancers 2 and 4, which are stimulated by dFOXO overexpression, was barely above background (Wei et al., 2016). Thus, predicting dFOXO responses from ChIP binding data is challenging. Similarly, we found that the presence of dFOXO-like binding motifs does not appear to correlate with with dFOXO response in S2 cells (data not shown). It is possible that our transient transfection assays, with dFOXO overexpression, may distort the true response of the enhancers to dFOXO; subsequent mutation of individual dFOXO-responsive elements within the endogenous gene, such as in 2a and 4ab, will be illuminating. Our studies did not conclusively determine whether the inhibitory effect of dFOXO on the T1 promoter and other enhancers is direct or indirect, but given the 72-hour timeline for the procedure, there would be time enough for known growth-inhibitory effects of dFOXO to broadly interfere with gene expression.

The use of reporter genes is a widespread and powerful tool for detailed assessment of regulatory regions. However, there are clear limitations to this reductionist approach when trying to gain an understanding of a large and complex regulatory locus. First, the demarcations of individual enhancers are arbitrary, carried out for efficiency in designing transfection assays. We show that many of the defined enhancers lose activity when divided into smaller fragments, thus there is every reason to expect that some of the divisions between the original 1.5 kbp elements fatally subdivided into a more complex element. We did however find that some elements, notably enhancer 2, were readily subdivided to find the dFOXO and EcR responsive portions. Furthermore, we were able to test the function of larger regions; by combining enhancer 2 and enhancer 3, we discovered that these enhancers appear to work additively, but with treatment of 20E, it appears that the combined region may be superstimulated, suggesting a synergistic action (Fig. 5B). On the endogenous locus, we imagine that the enhancers may work additively, or they may antagonize or synergize. Clearly, to establish this sort of function we will need to examine the function of the mostly intact, endogenous locus– a task that has become much more simplified in an era of CRISPR tools. However, in sum, the use of transfected reporters has allowed us to move considerably beyond our initial assessment, from defining elements that are necessary to elements that are sufficient for certain responses. Thus, this reductionist analysis of each enhancer can help us better understand the regulation of *InR*. Future work will focus on the endogenous locus, using diverse techniques to understand how these enhancers work as an ensemble. Delving into the mechanisms by which this gene is regulated will help us understand natural variation, providing insights into pathological states, development, and evolution.

## Materials & Methods

### Cloning

The *InR* introns were previously divided into ∼1.5 kbp fragments, the fragments that showed response to FOXO or 20E were broken down further into ∼300 bp and ∼600 bp (a-e and ab-de). Aside from Enhancer 22 which is longer than 1.5kb and is broken down into (22a-f, and 22ab-ef). The 300 bp and 600 bp fragments were cloned into the same luciferase reporter vector as previously described by Wei et al., 2016. The vector, p2T-Luc, is a luciferase reporter vector that also contains an ampicillin-resistant gene and a CMV-Rinella Promoter. Luciferase reporters were constructed previously by Ryu and Arnosti, 2003. The intronic fragments were cloned upstream of the T1 promoter between the Not1 and Asc1 sites of the vector and the transcription factor dFOXO was cloned into the pAX vector as previously described by Wei et al., 2016. All primers can be found in the table (Table 1) and were ordered from Integrated DNA Technologies, IDT (Owczarzy et al., 2008). All cloned fragments were checked for accuracy through Sanger Sequencing at Michigan State University Genomics Department.

### Transfections

Drosophila melanogaster S2 cells from the Drosophila Genomics Resource Center at the Indiana University of Bloomington were used for the cellular assays. 1.5 mL of 1-1.5 million/mL of cells were placed into each well of 6 well plates. 250 ng of the plasmids with the intronic fragments were expressed into the S2 cells with Effectene Transfection Reagent from Qiagen. For dFOXO, the cells were treated with either 200 ng dFOXO, 50 ng pBS or 114 ng pAX, and 136 ng pBS as the control. For 20E the cells were treated with either 250 ng pBS and 2 mL of the hormone ecdysone (10mg/mL) or 250 ng pBS and 2 mL of ethanol as the control. dFOXO transfections sat for 72 hours before being assayed. The 20E transfections sat for 24 hours with the expressed plasmids before adding 2 mL of ethanol or 20E, and then sat for 24 more hours and then assayed at 48 hours total.

### Luciferase Assays

Steady-Luc HTS Assay Kit from Biotium was used for the luciferase assays. Once transfections were ready to be assayed, the cells were spun down and resuspended in 230 mL Dulbecco’s Phosphate Buffered Saline. The resuspended cells are split into 3 technical replicates, 65 mL each, and pipetted into 96 well plates, then 65 mL of luciferin substrate is added to each well. Once luciferin is added the plate is left to sit for 10 minutes exactly before running on a Veritas Luminometer. Values from the luminometer are normalized to each of the corresponding full fragments’ basal activity. T1 was used as a negative control, and the full fragments were used as positive controls.

### Statistical Analysis

Luciferase assays for the 300 bp and 600 bp fragments were normalized to the full-size enhancer controls. We used Prism GraphPad Prism9 to perform multiple t-tests analysis of the control vs. treatment to check for statistical significance (GraphPad Prism version 9.0.0 MacOS, GraphPad Software, San Diego, California USA, www.graphpad.com).

### Sequence Analysis

We pulled 12 various species of Drosophila *InR* sequences from NCBI BLAST using the RefSeq genome sequence GCF_000001215.4 (Johnson et al., 2008). Sequences were aligned with Clustal Omega to analyze the conservation of the sequences throughout evolution (Sievers et al., 2011). Using the UCSC Genome Browser we were able to identify potential transcription factor binding motifs (Kent et al., 2002). Using MEME-Suite FIMO, we were able to upload the Drosophila alignment sequences and identify the specific loci of conserved transcription factor binding motifs of interest (Grant et al., 2011). JASPAR was used to find transcription factor binding motif sequence logos(Khan et al., 2018; Sandelin et al., 2004).

### Site-Directed Mutagenesis

Primers for site-directed mutagenesis were ordered from IDT and can be found in the primer chart mentioned above (Table 1) (Owczarzy et al., 2008). We used the Expand Long Enzyme PCR system from Millipore Sigma to make small deletions. After PCR, we used a Dpn1 restriction digest enzyme to digest any methylated DNA to obtain only the mutated plasmid we were interested in.

## Acknowledgments

We would like to thank the Department of Biochemistry and Molecular Biology and the Graduate School of Michigan State University for funding the Building Bridges Summer Internship to promote the recruitment of indigenous students. We are grateful to Ana-Maria Raicu and Sandhya Payankaulam for the mentoring and helpful discussions and the undergraduate student assistance from Carmen Cameron and Bhawna Vaswani. Subcloning of enhancer elements was initiated by Dr. Ali Bayram assisted by Gabby Hardy and Madeline Niblock.

